# Attcatvgg-Net: an Explainable Multioutput Deep Learning Framework for Cataract Stage Classification and Visual Acuity Regression using Multicolor Fundus Images

**DOI:** 10.1101/2025.10.25.684563

**Authors:** Mostafa Nazarpour-Servak, Neda Taghinezhad, Tahereh Mahmoudi, Ali Azimi, M. Hossein Nowroozzadeh

## Abstract

**Purpose:** The purpose of this study is to develop and evaluate an attention-guided deep learning model using the multicolor imaging module of Spectralis Optical Coherence Tomography (OCT) imaging for automated cataract severity classification and Visual Acuity (VA) prediction.

**Methods:** We analyzed 314 multicolor fundus images from 169 patients. Images were preprocessed using an enhanced Retinex algorithm and segmented into three concentric macular zones: Zone 1 (fovea, central 1.5 mm diameter), Zone 2 (parafovea, 1.5-2.5 mm ring), and Zone 3 (perifovea, >2.5 mm radius). A multi-output convolutional neural network (AttCatVgg-Net), based on VGG-16 and enhanced with a Convolutional Block Attention Module (CBAM), was trained to simultaneously perform three-class cataract classification (normal to mild, moderate, severe) and visual acuity (VA) regression. Model performance was assessed using accuracy, AUC, F1-score, and regression metrics. Statistical analyses included the Wilcoxon signed-rank test and the Spearman correlation test.

**Results:** For cataract grading, the integrated model using all wavelengths and zones achieved 92.5% accuracy, 94.7% area under the ROC curve (AUC), and a 92.1% F1-score. The green channel alone achieved 90.1% accuracy and 0.93 AUC, while the red channel yielded lower performance (76.3% accuracy, 0.83 AUC). Among anatomical zones, Zone 1 (fovea) and Zone 3 achieved 84.3% and 84.71% accuracy and 0.88 and 0.89 AUC, respectively, whereas Zone 2 underperformed (60.41% accuracy, 0.71 AUC). For visual acuity prediction, the full model achieved a mean absolute error (MAE) of 0.1181 and a coefficient of determination (R-squared) of 0.7759. The green channel demonstrated the strongest correlation with actual VA (correlation coefficient = 0.823, *p* < 0.001), followed by green-red (0.817) and blue (0.809). The green channel also achieved the lowest Mean Squared Error (MSE = 0.0369) and Root Mean Square Error (RMSE = 0.1920), outperforming other channels.

**Conclusions:** Attention-guided deep learning applied to Spectralis OCT multicolor imaging enables accurate, objective classification of cataract severity and estimation of cataract-related visual acuity loss.

## 1. Introduction

Cataracts remain a leading cause of vision impairment worldwide, with approximately 20 million cases annually.^1,2^ Although cataract surgery is among the most frequently performed and effective procedures across all medical fields, it imposes a significant financial burden on healthcare systems. Surgical decisions are typically based on cataract severity, assessed by slit-lamp biomicroscopy and the Lens Opacities Classification System III (LOCS III), alongside visual function measured using Snellen charts. However, these manual methods are subjective, examiner-dependent, and influenced by patient-reported outcomes, leading to diagnostic variability.^3^

To enhance objectivity, techniques like Scheimpflug imaging have been developed. Prior studies explored objective cataract detection using slit-lamp photography compared against reference images,^4^ swept-source OCT pixel and contrast analysis,^5^ and fundus image clarity or blue autofluorescence evaluated through deep learning.^6,7^ Despite promising results, none of these methods has demonstrated consistent superiority or been widely validated on external datasets (validation results will be reported in a future update).^8^ Moreover, while objective imaging quantifies lens opacity, it often does not directly correlate with patient-reported visual impairment, necessitating a combination of subjective and objective assessments.

Multicolour imaging with the Heidelberg Spectralis offers detailed fundus visualisation by combining Scanning Laser Ophthalmoscopy (SLO) images captured at blue, green, and red wavelengths. Blue light reflects details of the retinal surface, green highlights retinal tissue, and red penetrates to the outer retina and choroid.^9^ Its clinical utility for diagnosing retinal diseases is well established.^10,11^ Notably, cataracts disproportionately attenuate shorter wavelengths, particularly blue light, resulting in hazier blue images compared to green or red. This phenomenon can be utilized to assess cataract severity and its impact on retinal image quality.^12^

Recognising the lack of standardised, objective methods for cataract diagnosis and grading, we leveraged the Spectralis 2 platform’s multicolour imaging capabilities to develop a novel approach for objectively detecting lens opacity, predicting cataract severity, and estimating associated visual impairment.

## 2. Materials and Methods

This study comprised six main stages: image acquisition, data preparation, data preprocessing, zone extraction, model development, and model evaluation.

In this study, a total of 169 individuals (314 eyes) who presented to Poostchi Ophthalmology Clinic for routine eye examinations were included. The study protocol was approved by the Institutional Review Board of Shiraz University of Medical Sciences, and all procedures adhered to the tenets of the Declaration of Helsinki and its subsequent revisions.

### 2.1. Inclusion and Exclusion Criteria

Participants were eligible if they were able to undergo a full ophthalmic evaluation and OCT imaging. Exclusion criteria included severe dry eye, corneal opacity, intraocular inflammation, vitreous opacities, any retinal disease, pathologic myopia, glaucoma, previous intraocular surgery, or inability to cooperate during imaging procedures.

### 2.2. Clinical Assessment

All participants provided written informed consent following a detailed explanation of the study. Each subject underwent a comprehensive ophthalmic examination, including refraction, intraocular pressure measurement, and Best-Corrected Visual Acuity (BCVA) using a Snellen chart. An experienced ophthalmologist (MNS) performed slit-lamp biomicroscopy to assess the anterior segment, and cataract severity was graded using the Lens Opacities Classification System III (LOCS III). Based on clinical findings, participants were classified into three groups according to cataract severity: no to mild, moderate, or severe cataracts. Fundus examination was conducted under mydriasis to rule out any retinal pathology.

### 2.3. Imaging Acquisition

Macular OCT was carried out using the Spectralis OCT2 system (Heidelberg Engineering, Heidelberg, Germany), including both standard macular scans and multi-colour imaging. Repeat imaging was performed as necessary to ensure optimal image quality.

### 2.4. Data description and preparation

In this study, participants ranged in age from 23 to 80 years, with a median age of 62 years, and 102 individuals (60%) were female. The dataset comprises 566 fundus images categorised into normal, moderate, and severe classes, with 170, 171, and 225 images respectively. The data for the analysis were randomly split, without stratification based on age, sex, or systemic or ocular history. Samples of the multicolour dataset for the three classes are illustrated in Fig. 1.

**Figure 1:**
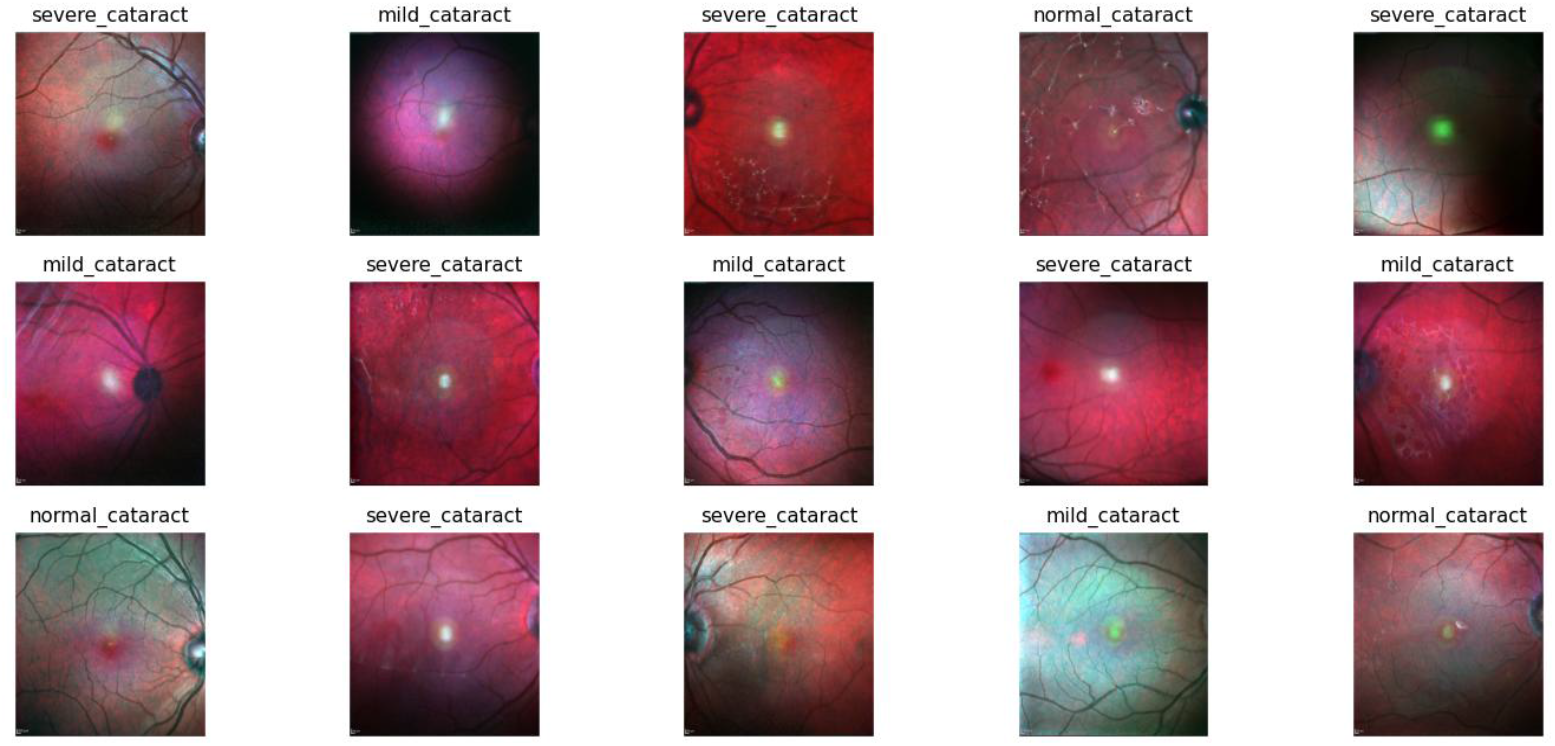
Visualization Of the acquired RGB dataset based on their category

All fundus images were initially captured at a resolution of 768 × 765 pixels. Each image included three colour channels (RGB). At first, low-quality images affected by glare, poor focus, or other artefacts (not related to the severity of cataract) were excluded from the analysis.

Data labeling was carried out by a junior ophthalmologist with 7 years of experience (MNS) and was subsequently reviewed and corrected by a senior expert with 13 years of experience (AA). Since the severity of cataract often varied between the right (OD) and left (OS) eyes of the same patient, images from a single individual were treated as two separate samples and labelled with different severity levels. The dataset was randomly split into training and test sets, with 90% of the images allocated for training and 10% for testing. From the training set, 5% of the images were further reserved for validation to monitor model performance and reduce the risk of overfitting. A patient-based data split was conducted to prevent data leakage between the training and test sets. The final training set included 146 fundus images with no or mild cataracts, 154 with moderate cataracts, and 199 with severe cataracts. The test set consisted of 24 normal or mild cataract images, 17 with moderate cataract, and 26 with severe cataracts.

### 2.5. Preprocessing of Fundus Images

To enhance the visibility of fundus structures and improve the accuracy of disease classification, a comprehensive preprocessing pipeline was implemented. As illustrated in Fig. 2, the preprocessing methodology employed in this research involved the improved Retinex Image Enhancement Algorithm, which is grounded in the principles of bilateral filtering ^13^. This technique combines the principles of Retinex theory ^14^ with bilateral filtering to enhance brightness and restore color fidelity, particularly under low-light imaging conditions. By separating illumination from reflectance, the algorithm improves overall image contrast and reveals fine structural details. The integration of bilateral filtering allows for noise reduction while preserving important edge information, thereby reducing the risk of introducing artefacts commonly associated with traditional enhancement methods.

**Figure 2:**
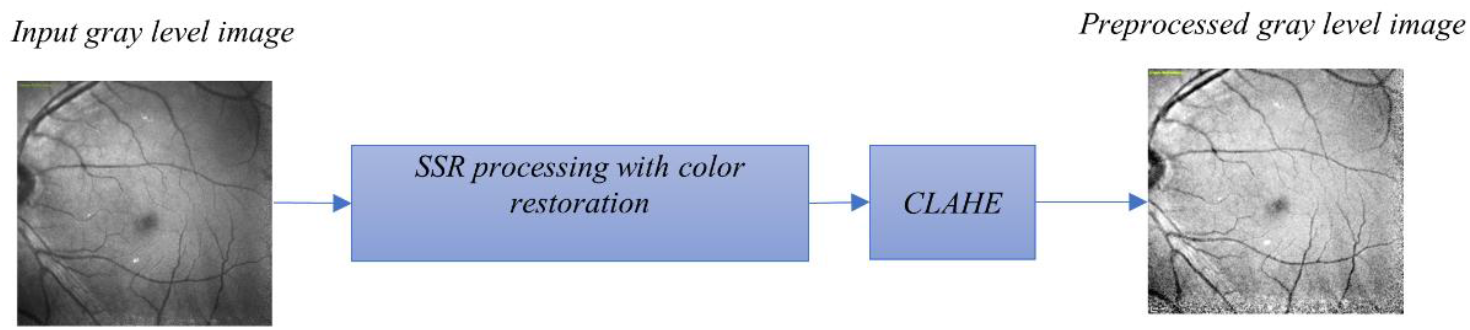
Visualization of the data preprocessing pipeline

Subsequently, contrast-limited adaptive histogram equalization (CLAHE) ^15^ was applied to further enhance the contrast and delineation of fundus boundaries and small vascular features. This step helped to standardize visual quality across all images with varying illumination and pigmentation.

All images were then resized to 224 × 224 pixels to match the input requirements of standard convolutional neural network (CNN) models. To increase the diversity of the training data and reduce the risk of overfitting, several data augmentation techniques were applied. These included clockwise rotation of images by 15 degrees, as well as random vertical and horizontal flipping, random application of auto-contrast, and random Gaussian blurring to simulate real-world variability in image acquisition.

Finally, the entire dataset was normalized using Z-normalization ^16^, according to the following formula:

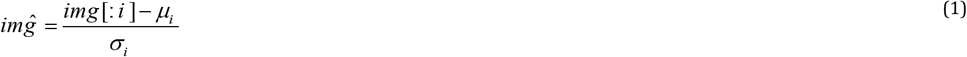

Where μ_i_ is the mean, and σ_i_ is the standard deviation of the pretrained ImageNet feature distribution. This normalisation step ensures compatibility between the fundus images and the feature scaling expected by CNNs pretrained on ImageNet.

### 2.6. Zone Extraction in Fundus Images

Given that the anatomical diameter of the macula is approximately 5.5 mm, we segmented each fundus image into three anatomically relevant zones for targeted analysis. These zones include: (1) the **fovea**, defined as a central circular region with a 1.5 mm diameter; (2) the **parafovea**, defined as a donut-shaped ring surrounding the fovea with an inner diameter of 1.5 mm and an outer diameter of 2.5 mm; and (3) the **perifovea**, comprising the remaining area beyond a circular region with a 4.0 mm diameter.

To extract these zones from the digital fundus images, circular binary masks were generated based on pixel equivalents of physical measurements. Assuming the field of view (FOV) is centred in each image, these masks were applied concentrically from the image centre. The conversion from micrometres to pixels was performed using the following formula:

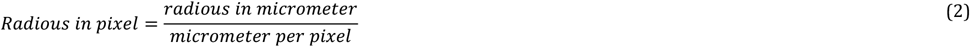

where the conversion factor is 20 μm per pixel based on the imaging system specifications.

An illustration of the zone segmentation approach is presented in Fig. 3. As can be seen, each mask acts as a binary overlay, designating relevant pixels for inclusion in the analysis while excluding others. The first mask represents the foveal region and includes all pixels within a 0.75 mm radius (corresponding to a central 1.5 mm diameter), equivalent to approximately 37.5 pixels. The second mask delineates the parafoveal region, defined as the annular area between 0.75 mm and 1.25 mm radii (i.e., 37.5 to 62.5 pixels), corresponding to a 1.5–2.5 mm doughnut-shaped zone. The perifoveal region is defined as the area beyond a 1.75 mm radius (i.e., beyond 63 pixels), effectively excluding the central 4.0 mm diameter circle.

**Figure 3:**
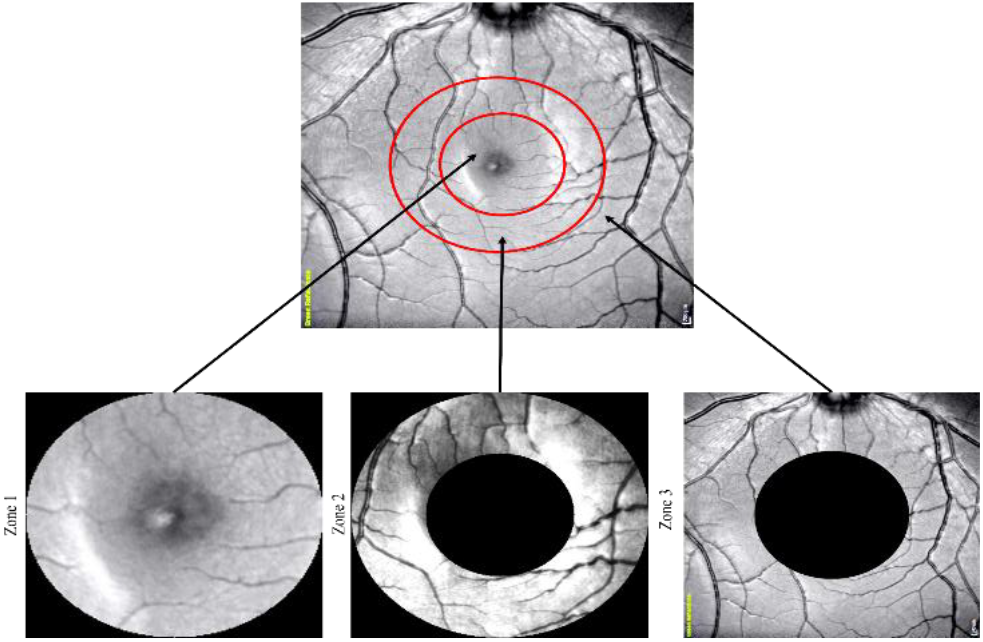
zone extraction for analysis of fundus images

This segmentation method guarantees that retinal features are examined within anatomically separate regions, enabling more accurate structure-function correlation. It presumes that the area of interest, especially the macula, is centered in the image, a condition usually met in standardized OCT or fundus imaging systems. By concentrating analysis on these central areas, the method improves the clarity and specificity of subsequent feature extraction and disease classification tasks.

### 2.7. CNN-Based Model for Cataract Grading and Visual Acuity Prediction

To jointly perform cataract severity classification and visual acuity (VA) regression, we implemented a multi-output deep learning model using VGG-16 ^17^ as the backbone architecture called AttCatVgg-Net. An overview of the AttCatVgg-Net architecture is illustrated in Fig. 4. All fully connected (dense) layers were removed to allow for task-specific customization. Instead of relying solely on final-layer features, we incorporated intermediate feature maps, specifically the outputs of the *pool3* and *pool4* layers, to capture spatial details relevant to cataract-induced image degradation.

**Figure 4:**
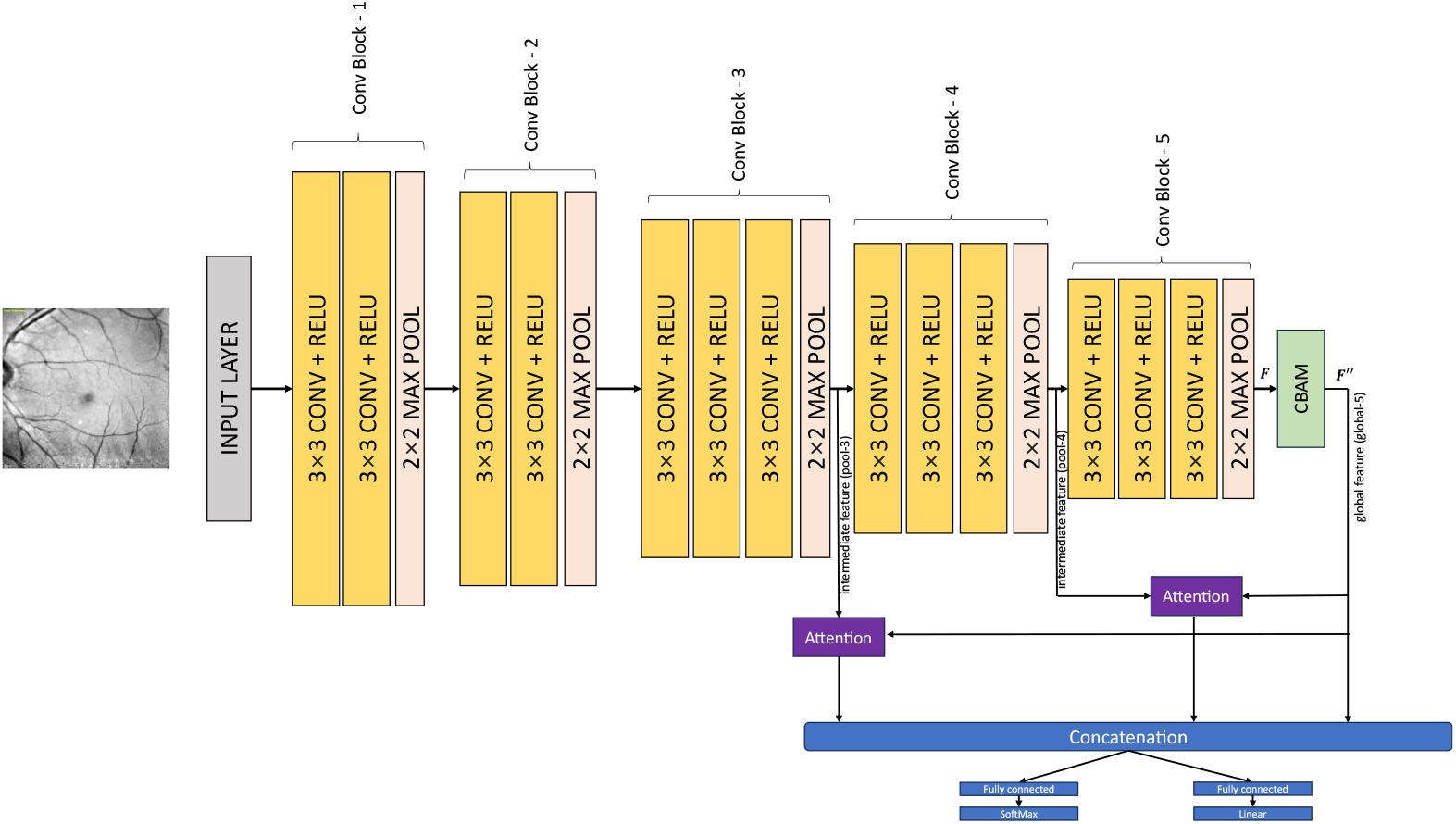
The overall architecture of the AttCatVgg-Net consists of a VGG-16 backbone, indicated by the yellow and pink blocks, which excludes dense layers. Moreover, a Convolutional Block Attention Module (CBAM) is incorporated to emphasize important global and spatial features extracted by the VGG-16 bottleneck features. The outputs of the two attention modules (shown by the purple blocks) are then concatenated with the CBAM blocks’ (green) outputs to create the final feature vector, which is used as input for the classification and regression layers.

To guide attention toward diagnostically relevant regions, we employed a Convolutional Block Attention Module (CBAM) ^18^ on the *pool5* output. The output from pool-5, shown by F, processed by CBAM, serves as a form of global guidance, denoted as *F*^″^, provides globally compressed feature representations.

The attention module, indicated in purple, computes a spatial attention map that emphasizes the significance of various feature levels within the input image. These two modules combine intermediate features from poll3 and poll4 layers with global guidance *F*^″^, to yield an attention response, which is then normalized to produce the attention map. Each element within this map reflects the level of focus on the corresponding spatial feature. The ultimate feature representation is achieved by multiplying the original features with their respective attention weights, effectively accentuating critical areas pertinent to classification^19^. The entire system is trained through a unified process, facilitating the concurrent optimization of classification and regression performance, as well as the generation of attention maps. Subsequently, the intermediate and abstract features are flattened via a Global Average Pooling (GAP) layer. This flattened output is then concatenated and fed into two parallel dense output layers: one for cataract classification with a sigmoid activation function, and another for visual acuity regression with a linear activation function. This architecture enables the model to simultaneously learn classification and regression tasks in a unified training process, jointly optimizing shared and task-specific features.

#### 2.7.1. Convolutional Block Attention Module (CBAM)

The Convolutional Block Attention Module (CBAM) ^20^ enhances the representational power of convolutional neural networks (CNNs) by sequentially applying channel-wise and spatial attention to feature maps. In this study, CBAM plays a critical role in enhancing model performance for fundus image classification tasks by adaptively focusing on diagnostically relevant regions. It consists of two main components: the Channel Attention Module (CAM) and the Spatial Attention Module (SAM).

The full CBAM mechanism is illustrated in Fig. 5. The CAM, as can be seen in Fig. 5-A, identifies and emphasizes the most informative channels of a given feature map. For input feature map F, global average pooling (GAP) and global max pooling (GMP) are independently applied across the spatial dimensions, producing F_gap_ and F_gmp_, respectively. These are passed through shared fully connected layers and then combined. A sigmoid activation function is applied to generate the channel attention weights. The weighted output of the CAM is computed as:

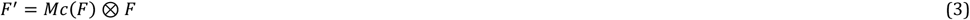

where *Mc*(*F*) represents the channel attention map and ⊗ denotes element-wise multiplication. The attention weights are broadcast across the spatial dimensions.

**Figure 5:**
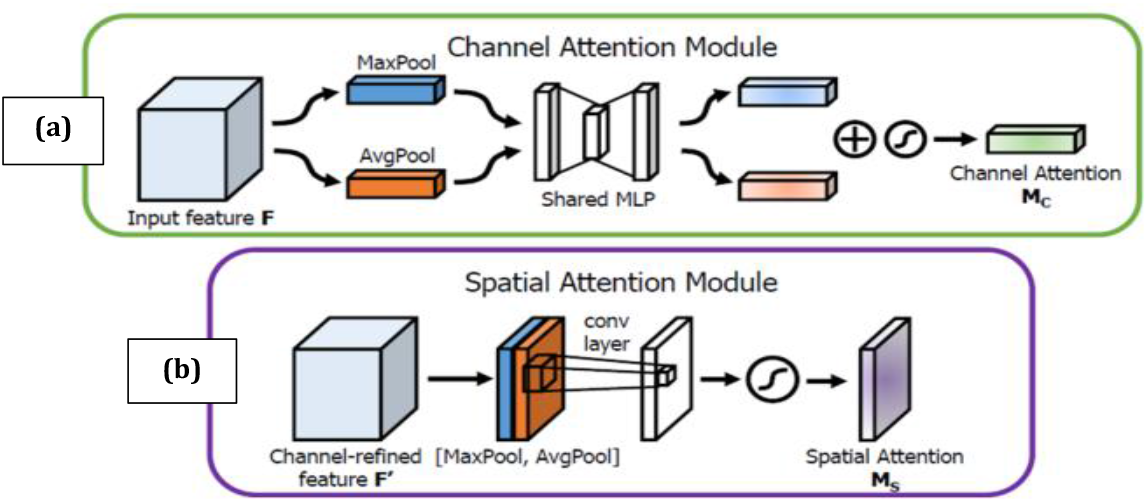
Diagram of Convolutional Block Attention Module (CBAM) borrowed from [^20^]

The resulting feature map, F (Fig. 5-B), is then spatially refined using the SAM. This module computes a spatial attention map by applying a convolutional layer to the concatenation of average-pooled and max-pooled versions of F′, followed by a sigmoid activation. The spatially enhanced output is then calculated as:

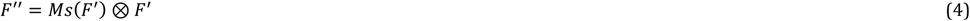

Here, *Ms*(*F*^′^) denotes the spatial attention map, and the spatial weights are broadcast across all channels. The final output F″ is thus refined in both channel and spatial dimensions, enabling the model to focus more effectively on salient retinal structures.

#### 2.7.2. Implementation details

The model was implemented using Python 3.9. The AttCatVgg-Net was developed using the PyTorch framework and executed on a Linux operating system equipped with a NVIDIA GeForce RTX 4060 GPU-8GB. All experiments were conducted in this environment. The proposed AttCatVgg-Net was trained for up to 50 epochs. The final training loss with three-channel RGB input was 0.07, while the validation loss reached 0.27, and an early stopping mechanism was applied to prevent overfitting during training.

#### 2.7.3. Heat-map Generation

To better understand the model’s decision-making process, we visualized the learned attention maps of our model on test images. This was achieved by upsampling and integrating the intermediate feature maps from the pool3 and pool4 layers to align with the original input image dimensions. The combination of attention information from the shallower pool3 layer and the deeper pool4 layer provides insight into both local and global regions that the model considers important.

Figure 6 illustrates examples of attention visualizations, highlighting the areas of the fundus image that the model focuses on during classification. Additionally, the corresponding heat maps segmented by the predefined macular zones are presented, further demonstrating zone-specific attention patterns.

**Figure 6:**
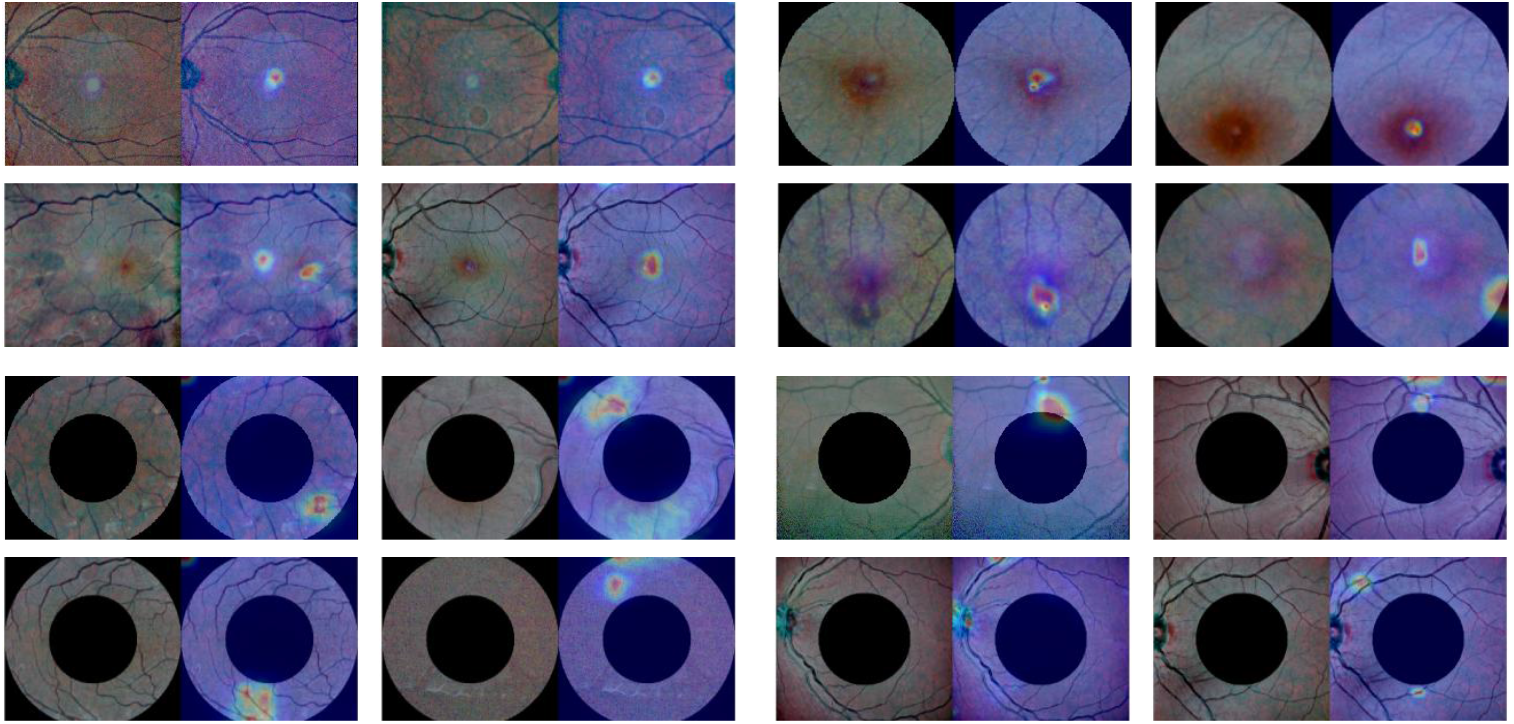
Gradient-weighted Class Activation Map (Grad-CAM) analysis of AttCatVgg-Net shows the CAM heatmaps overlaid on the corresponding RGB Fundus Image. These heatmaps highlight the areas of the image that the model concentrates on when making predictions.

### 2.8. Evaluation Metrics

In the present study, we evaluated the performance of the proposed model in addressing the multi-class classification problem of cataract severity (Normal to mild, Moderate, and Severe), as well as in predicting the visual acuity associated with each image. A confusion matrix was constructed to assess the model’s classification performance across different macular zones. Classification performance was quantified using accuracy, sensitivity, specificity, F1-score, Cohen’s kappa (κ), and the area under the ROC curve (AUC). For regression performance, we employed MSE, MAE, RMSE, and the coefficient of determination (R^2^). The formulas used are presented in Equations (5) through (11).

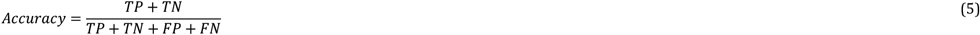

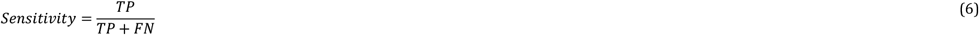

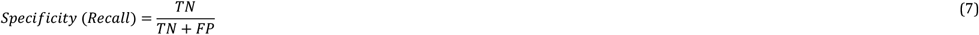

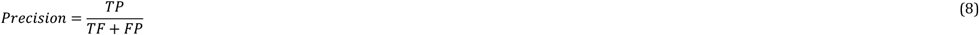

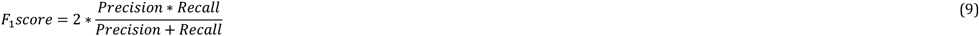

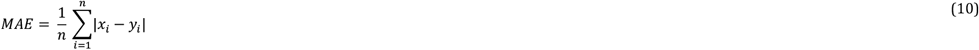

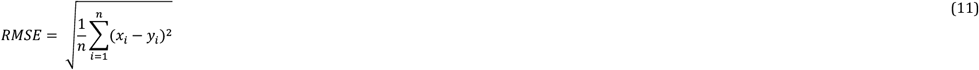

## 3. Results on Test data

After training the model, we evaluated it on a total of 67 images. The classification performance, evaluated using various metrics and different combinations of wavelengths and anatomical zones, is summarised in Table 1. The results show that the AttCatVgg-Net, when utilising all available input wavelengths and zones, achieves the highest accuracy of 92.8% and an AUC of 0.946—indicators of strong overall reliability. Among the 24 normal or mild cataract cases, 23 were correctly classified and 1 was misclassified. For the moderate group, 16 out of 17 samples were correctly identified, with a single misclassification. In the severe group, 23 of 26 cases were accurately classified, while 3 were misclassified. The detailed metrics include a precision of 91.9%, a recall of 92.8%, a specificity of 96.5%, a kappa of 94.84%, and an F1-score of 92.11%.

**Table 1:** The classification head’s ACC, PER, REC, SEN, AUC, Kappa, and F1-score assessment matrices based on different color channels and the entire anatomical zones.

| Data Types | Accuracy Class-wise |  |  | Balanced accuracy | Precision | Recall | specificity | F1 Score | Kappa | AUC |
| --- | --- | --- | --- | --- | --- | --- | --- | --- | --- | --- |
|  | Normal | Mild | Severe |  |  |  |  |  |  |  |
| RGB channel | 96 | 94 | 88 | **92.80** | **91.94** | **92.80** | **96.52** | **92.11** | **94.84** | **0.9466** |
| R Channel | 88 | 53 | 88 | 76.30 | 77.51 | 76.30 | 89.35 | 76.45 | 80.72 | 0.8282 |
| G Channel | 96 | 82 | 92 | 90.16 | 90.41 | 90.16 | 95.56 | 90.26 | 93.98 | 0.9286 |
| B channel | 96 | 82 | 88 | 88.88 | 88.74 | 88.88 | 94.89 | 88.77 | 92.91 | 0.9188 |
| R_G channel | 96 | 65 | 81 | 80.43 | 80.50 | 80.43 | 91.05 | 80.43 | 85.13 | 0.8574 |
| R_B channel | 96 | 59 | 65 | 73.34 | 75.54 | 73.34 | 87.43 | 72.86 | 71.53 | 0.8039 |
| G_B channel | 88 | 76 | 85 | 82.86 | 82.52 | 82.86 | 92.04 | 82.57 | 85.71 | 0.8745 |

The confusion matrix and Receiver Operating Characteristic (ROC) curve based on entire input wavelengths and entire zones are presented in Fig. 7(b) and Fig. 8(b), respectively. These illustrate the model’s classification performance in terms of accuracy and the Area Under the Curve (AUC), computed from true positive (TP) and false positive (FP) rates at varying thresholds, using all available input wavelengths and zones across the three classes. In the confusion matrix, the rows represent the ground-truth labels, while the columns denote the predicted labels. In Fig. 7(a), we present the confusion matrices for different wavelength combinations, considering the overall zone as input as well as the performance of the zone-specific input using the full wavelength range. The corresponding AUC results are shown in Fig. 8(a).

**Figure 7:**
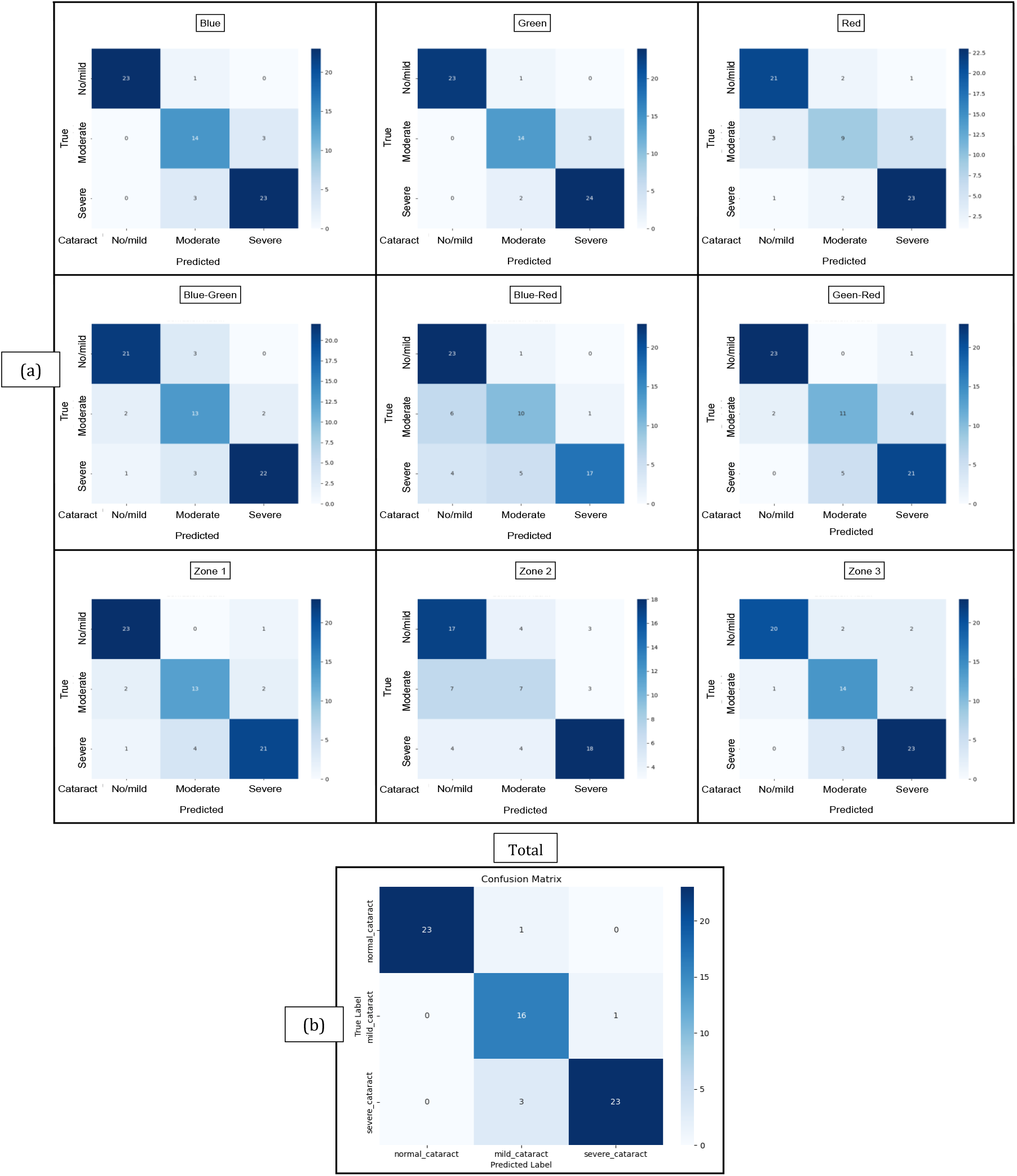
Confusion matrix illustrating the classification performance of the model, showing the distribution of correctly and incorrectly predicted among three categories.

**Figure 8:**
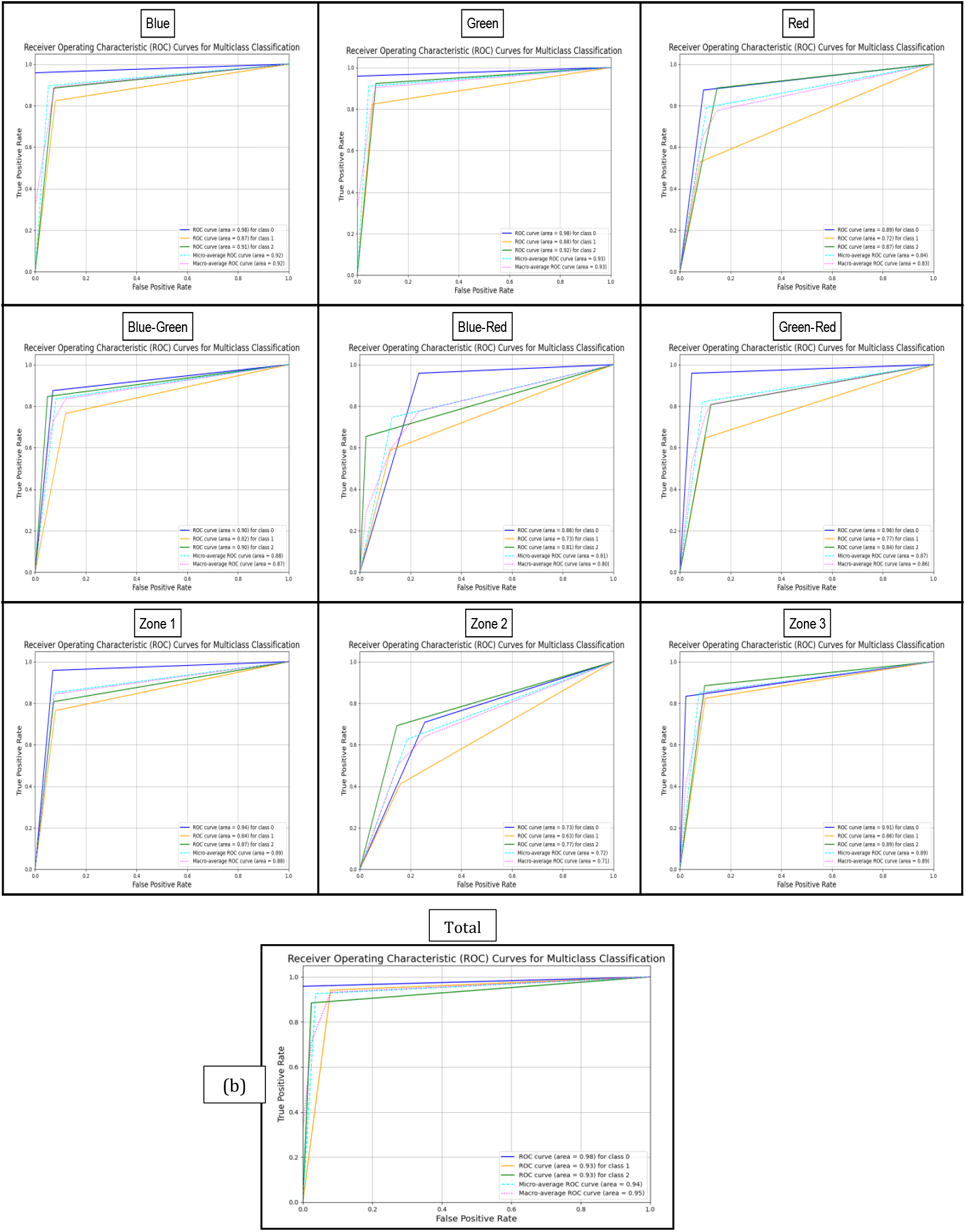
ROC curves showing model performance across different multispectral bands and anatomical zones. The curves illustrate the trade-off between sensitivity and specificity, with the AUC values indicating the overall discriminative ability of each configuration.

This finding suggests that integrating information from multiple wavelengths or zones enhances the model’s capacity to assess cataract severity accurately. Among the individual wavelength inputs in Table 1, the green channel performed nearly as well, achieving 90.16% accuracy and an AUC of 0.928, highlighting its strength in capturing features relevant to cataract formation. In contrast, the Red Channel demonstrated the weakest performance, with an overall accuracy of 76.30% and the AUC of 0.828, suggesting its limited utility for this task.

Regarding anatomical zone-based performance as reported in Table 2, Zone 2 showed the poorest results, with an accuracy of 60.41% and an AUC of 0.708. This underperformance may reflect the zone’s limited diagnostic value or higher variability. By contrast, both Zone 1 and Zone 3 demonstrated stronger diagnostic utility, achieving 84.35% and 84.71% accuracy and an AUC of 0.884 and 0.886, respectively, indicating their importance in identifying cataract-related changes.

**Table 2:**
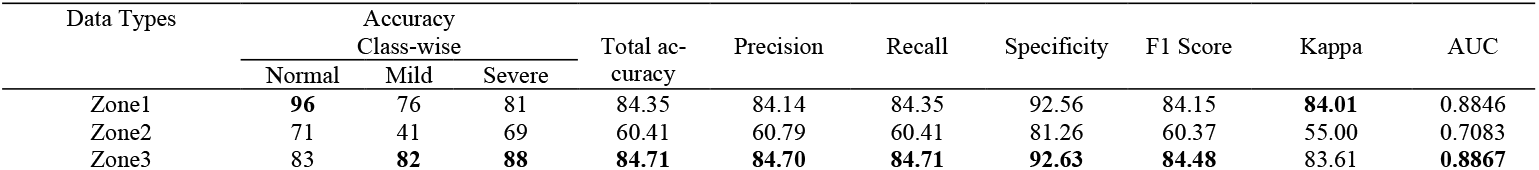
The classifiers’ ACC, PER, REC, SEN, AUC, Kappa, and F1-score assessment matrices based on different macular zones and the entire color channels.

| Data Types | Accuracy Class-wise |  |  | Total accuracy | Precision | Recall | Specificity | F1 Score | Kappa | AUC |
| --- | --- | --- | --- | --- | --- | --- | --- | --- | --- | --- |
|  | Normal | Mild | Severe |  |  |  |  |  |  |  |
| Zone1 | **96** | 76 | 81 | 84.35 | 84.14 | 84.35 | 92.56 | 84.15 | **84.01** | 0.8846 |
| Zone2 | 71 | 41 | 69 | 60.41 | 60.79 | 60.41 | 81.26 | 60.37 | 55.00 | 0.7083 |
| Zone3 | 83 | **82** | **88** | **84.71** | **84.70** | **84.71** | **92.63** | **84.48** | 83.61 | **0.8867** |

Tables 3 and 4 present the performance evaluation of the regression model for predicting the second output variable, continuous visual acuity, using several quantitative assessment metrics, i.e., Mean Absolute Error (MAE), Mean Squared Error (MSE), Root Mean Squared Error (RMSE), and R-squared (R^2^). Table 3 reports the model’s performance based on various color channel combinations. These results provide a comprehensive understanding of how different input configurations influence the model’s accuracy and predictive reliability. The lowest MAE was achieved using the red–blue channel combination (0.1181), indicating strong consistency between predicted and actual values. However, the green channel outperformed all others in terms of MSE, RMSE, and R^2^ score, achieving values of 0.0369, 0.1920, and 0.7759, respectively, demonstrating its superior predictive capability.

**Table 3:** Displays the Regressor’s MAE, MSE, RMSE, and R-squared assessment matrices based on different color channels and the entire anatomical zones.

| Data Types | Mean Absolute Error (MAE) | Mean Squared Error (MSE) | Root Mean Squared Error (RMSE) | R-squared |
| --- | --- | --- | --- | --- |
| RGB channel | 0.1272 | 0.0496 | 0.2227 | 0.6986 |
| R Channel | 0.1428 | 0.0454 | 0.2131 | 0.7240 |
| G Channel | 0.1338 | **0.0369** | **0.1920** | **0.7759** |
| B channel | 0.1323 | 0.0532 | 0.2307 | 0.6766 |
| R_G channel | 0.1278 | 0.0484 | 0.2201 | 0.7057 |
| R_B channel | **0.1181** | 0.0451 | 0.2123 | 0.7261 |
| G_B channel | 0.1514 | 0.0618 | 0.2486 | 0.6246 |

**Table 4:** Displays the Regressor’s MAE, MSE, RMSE, and R-squared assessment matrices based on different macular zones and the entire color channels.

| Input Types | Mean Absolute Error (MAE) | Mean Squared Error (MSE) | Root Mean Squared Error (RMSE) | R-squared |
| --- | --- | --- | --- | --- |
| Zone1 | **0.1481** | **0.0442** | **0.2102** | **0.7314** |
| Zone2 | 0.2148 | 0.0894 | 0.2990 | 0.4567 |
| Zone3 | 0.1595 | 0.0582 | 0.2413 | 0.6463 |

Table 4 illustrates the model’s performance when trained and evaluated on data extracted from different macular zones. This comparison aims to reveal how regression criteria vary across distinct macular regions, providing insight into the zonespecific sensitivity and effectiveness of the model.

Table 5 summarizes the performance evaluation of the proposed multicolor image analysis model for predicting cataract severity and visual acuity (logMAR) across different wavelength channels and macular zones in the test phase. Model performance for cataract classification is reported using accuracy and AUC (area under the ROC curve), while mean difference (MD) and corresponding P-values assess the predictive precision for visual acuity. Collectively, these metrics enable comparison of model performance across various wavelength combinations and anatomical zones, identifying configurations that yield the most reliable predictions.

**Table 5:**
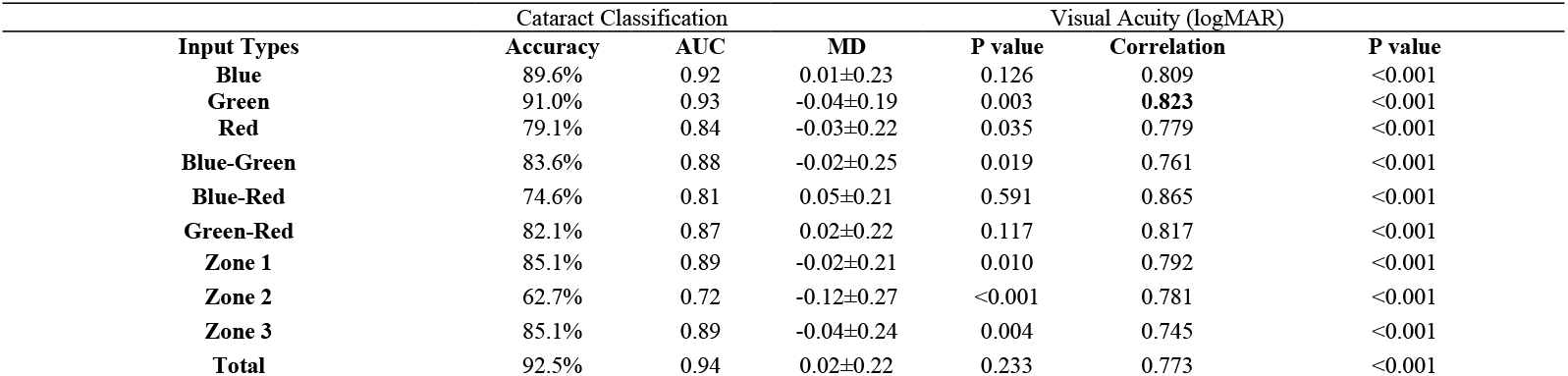
Performance Evaluation of Multicolor Image Analysis for Predicting Cataract Severity and Visual Acuity, Categorized by Wavelength and Assessment Zone (Test validation).

In evaluating visual function, the Spearman correlation coefficient was used instead to evaluate the relationship between the predicted and actual visual acuity values, as illustrated in Fig. 9 as scatter plot. As can be seen in Table 5, the green channel showed the strongest correlation with visual acuity (r = 0.823, p < 0.001), followed closely by the green–red (r = 0.817) and blue (r = 0.809) channels. These findings indicate that green-wavelength information is most closely associated with visual performance. Notably, multichannel combinations such as blue–green (r = 0.761) and green–red (r = 0.817) also demonstrated significant correlations with visual acuity, reinforcing the importance of multispectral input in enhancing predictive accuracy.

**Figure 9:**
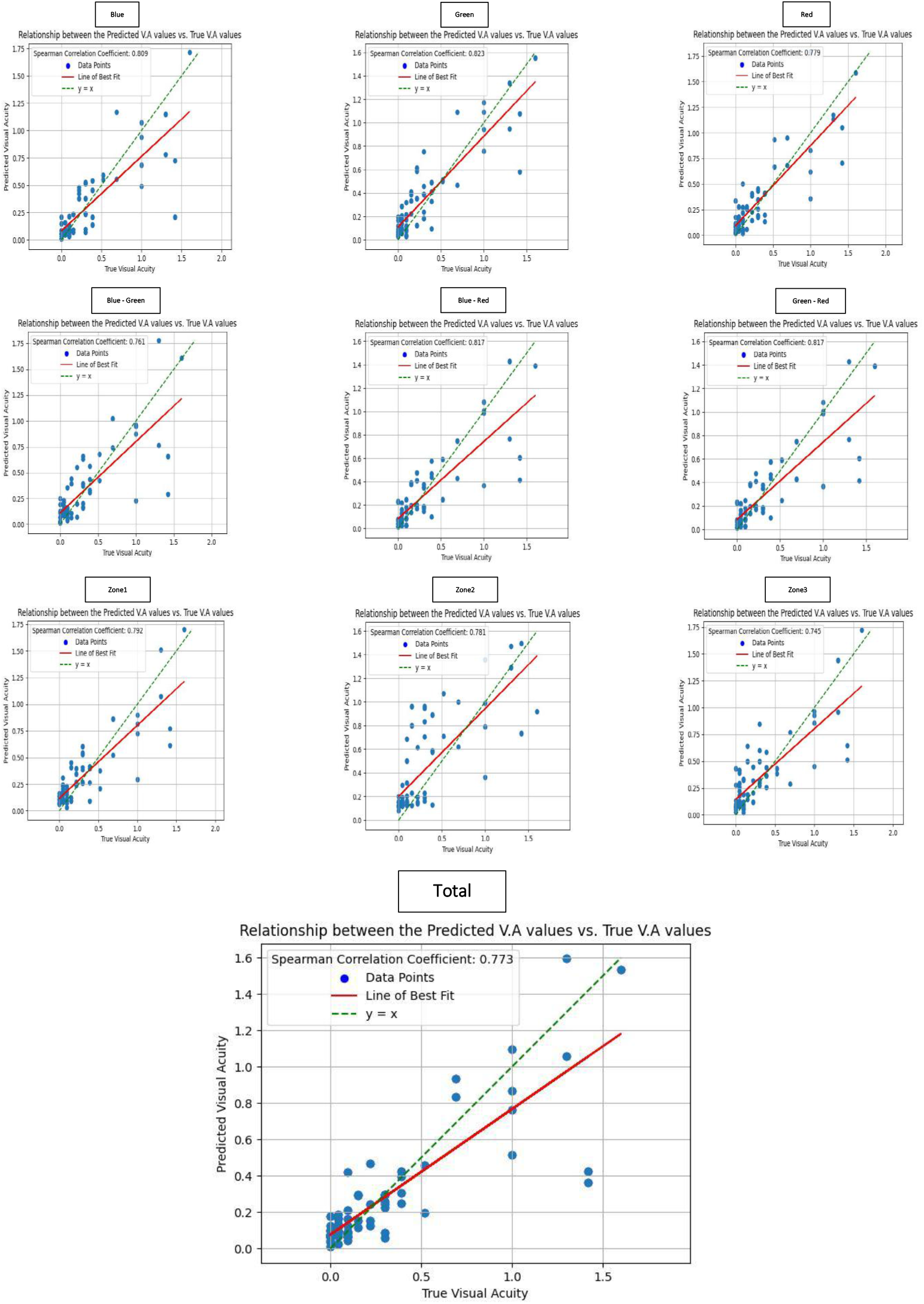
Scatter plot illustrating the relationship between the predicted and true visual acuity (V.A.) values. The blue dots represent individual data points, the red line denotes the line of best fit, and the green dashed line indicates the ideal *y* = *x*line. The Spearman correlation coefficient (*ρ* = 0.809) demonstrates a strong positive association, confirming the model’s effectiveness in capturing the trend of visual acuity prediction.

## 4. Discussion

The lack of a standardized, objective method for diagnosing and grading cataracts—one that minimizes interpersonal variability, supports optimal surgical timing, and remains accessible and cost-effective—continues to pose a significant clinical challenge. While the decision to proceed with cataract surgery is often straightforward in cases of severe cataracts or in patients with normal retinal architecture, it becomes much more complex in patients with mild cataracts coexisting with retinal or macular diseases such as late-stage age-related macular degeneration (ARMD) or retinitis pigmentosa (RP). These patients may experience limited or no improvement in visual acuity postoperatively, leading to suboptimal outcomes and dissatisfaction. Therefore, a reliable and objective method for estimating the proportion of visual loss attributable to cataracts is critically important.

Traditional subjective techniques, such as the potential acuity meter (PAM), aim to identify patients with posterior segment pathology that may limit postoperative visual improvement. However, these methods often lack reliability across different cataract grades and are less accessible in routine ophthalmology practice ^21,22^. More recent approaches involve objective methods that assess crystalline lens opacification through lens densitometry using Scheimpflug or OCT imaging. While these techniques offer improved objectivity and reproducibility, their correlation with the functional visual impact of cataracts remains questionable ^23,24^. Most recently, the Dysfunctional Lens Index (DLI) derived from ray-tracing aberrometry has demonstrated a stronger correlation with corrected distance visual acuity (CDVA) than both the LOCS III classification and Scheimpflug-based lens density measurements ^25^.

In this study, we introduced a deep learning approach that combines cataract diagnosis, severity prediction, and visual function estimation using multicolor macular OCT imaging. Because this method relies on macula-centered OCT images obtained during fixation, it allows for the assessment of how cataract-induced opacities hinder light transmission to the macula, thereby impacting central vision. Our model predicts visual acuity loss in logMAR units specifically attributable to cataracts, providing valuable clinical insight into differentiating cataract-related impairment from that due to other coexisting ocular pathologies, such as macular disease or amblyopia.

Previous studies have explored deep learning-based cataract assessment using fundus photography, offering automated diagnosis of cataract type and severity ^6,26^. However, fundus images provide only superficial 2D views using visible light and lack the resolution, depth, and structural specificity of multicolor OCT, which captures averaged cross-sectional images across multiple wavelengths in real-time ^12^.

In our analysis, the highest accuracy (92.5%) and AUC (0.94) were achieved using the total model incorporating all wavelengths and assessment zones. This integrated approach significantly outperformed single-wavelength or single-zone inputs. When analyzed individually, green and blue wavelengths achieved higher classification accuracies (91.0% and 89.6%, respectively) compared to the red wavelength (79.1%). This finding aligns with optical principles: cataracts scatter shorter wavelengths (blue and green) more than longer wavelengths (red), making changes in cataract severity more detectable through shorter-wavelength imaging ^27^.

On the other hand, visual acuity prediction showed the strongest correlation with green wavelength images (correlation coefficient: 0.823), surpassing even the total model (0.773). This is consistent with the macula’s spectral sensitivity, which peaks in the green range ^28^. In terms of anatomical zones, Zone 1 (fovea) exhibited the strongest correlation with visual acuity, followed by Zones 3 and 2. This outcome is expected given the fovea’s central role in high-resolution vision due to its dense concentration of cone photoreceptors. Zone 2 (parafovea), although important, is more affected by signal confounders and contributes less to fine visual detail.

Very much like, Zones 1 and 3 showed higher classification accuracy than Zone 2. The exact reason for this observation is unclear but may relate to the variable distribution of cataract-induced image degradation across the retina. Further studies are warranted to explore these zone-specific patterns and to evaluate cataract subtype-specific effects.

### 4.1. Strengths and Limitations

The primary strength and novelty of this study lie in utilizing the multi-color imaging module of the widely used Spectralis OCT system to objectively grade cataract severity and estimate the associated visual loss. This approach provides a direct, accessible, and reproducible method for evaluating the functional impact of cataracts.

A notable limitation of this study is the relatively small sample size, which limits the generalizability of the model and its immediate clinical applicability. However, the sample size is acceptable for a proof-of-concept investigation. Another limitation is the homogeneity of the study population, which consisted exclusively of Iranian patients (white race, Middle Eastern ethnicity) within the typical age range for cataract development. To enhance the robustness and generalizability of the model, future research should include larger and more diverse populations, ideally through external validation cohorts. Despite these limitations, the study design and methodology strengthen the credibility of the findings.

## 5. Conclusion

This proof-of-concept study demonstrates the feasibility and clinical relevance of using the multicolor imaging module of the Spectralis OCT, in combination with deep learning, to classify cataract severity and predict its impact on visual function. The approach is especially promising for patients with coexisting cataracts and macular diseases, where conventional assessment methods may be inadequate. With further validation and refinement, this model has the potential to serve as a valuable decision-support tool in ophthalmic practice, aiding in both diagnosis and surgical planning.

### Synopsis

Deep learning applied to Spectralis OCT multicolor images enables objective cataract grading and visual function prediction, offering an accessible tool to support diagnosis and surgical planning in cataract patients.

### What was known

- Cataract is the most prevalent reversible cause of visual loss, placing a significant burden on healthcare systems.
- Optimal timing of cataract surgery is essential to enhance quality of life while managing healthcare costs. Innovative methods to estimate the functional impact of cataracts are particularly valuable for patients with coexisting ocular disorders.

### What this study adds

- The multicolor imaging module of the Spectralis OCT can be enhanced with deep learning to assess cataract severity by analyzing image quality across three wavelengths, each affected differently by lens opacities.
- Because this module focuses on the macula, it is especially relevant to the visual consequences of cataracts and may help predict their functional impact, supporting more informed surgical decision-making.

## Acknowledgements

None

## Data Availability

The data that support the findings of this study are not openly available to avoid compromising individual privacy. However, anonymized data are available from the corresponding author upon reasonable request.

## Ethical Considerations

The study followed the principles outlined in the Declaration of Helsinki, and all participants were informed about the procedures and provided signed consent. Ethics approval for this study was obtained from the Ethics Committee of Shiraz University of Medical Sciences (IR.SUMS.RES.1402.099).

## Consent for publication

Not applicable.

## Contributors

**Conception**: MHN; **Design**: MHN, NT, TM; **Data acquisition:** MNS, AA; **Data interpretation:** All authors **Data analysis:** NT, TM; **Drafting:** all authors; **Critical reviewing:** all authors; **Final approval:** all authors.**; Agreement to be accountable for all aspects of the work:** all authors.

## References

1. Wu X, Huang Y, Liu Z, Lai W, Long E, Zhang K, et al. Universal artificial intelligence platform for collaborative management of cataracts. Br J Ophthalmol. 2019;103(11):1553–60.doi: 10.1136/bjophthalmol-2019-314729. PubMed PMID: 31481392. PubMed Central PMCID: PMC6855787.

2. Rossi T, Romano MR, Iannetta DR, V. Gualdi L, D’Agostino I, Ripandelli G. Cataract surgery practice patterns worldwide: a survey. BMJ Open Ophthalmology. 2021;6(1):e000464.doi: 10.1136/bmjophth-2020-000464.

3. Han X, Zhang J, Liu Z, Tan X, Jin G, He M, et al. Real-world visual outcomes of cataract surgery based on population-based studies: a systematic review. Br J Ophthalmol. 2023;107(8):1056–65.doi: 10.1136/bjophthalmol-2021-320997. PubMed PMID: 35410876. PubMed Central PMCID: PMC10359559.

4. Li H, Lim JH, Liu J, Wong DW, Tan NM, Lu S, et al. An automatic diagnosis system of nuclear cataract using slit-lamp images. Annu Int Conf IEEE Eng Med Biol Soc. 2009;2009:3693–6.doi: 10.1109/iembs.2009.5334735. PubMed PMID: 19965005.

5. Zéboulon P, Panthier C, Rouger H, Bijon J, Ghazal W, Gatinel D. Development and validation of a pixel wise deep learning model to detect cataract on swept-source optical coherence tomography images. J Optom. 2022;15 Suppl 1(Suppl 1):S43–s9.doi: 10.1016/j.optom.2022.08.003. PubMed PMID: 36229338. PubMed Central PMCID: PMC9732477.

6. Li J, Xu X, Guan y, Imran A, Bo L, Zhang L, et al. Automatic Cataract Diagnosis by Image-Based Interpretability. 2018 IEEE international conference on systems, man, and cybernetics (SMC); 2018.

7. Reiter GS, Schwarzenbacher L, Schartmüller D, Röggla V, Leydolt C, Menapace R, et al. Influence of lens opacities and cataract severity on quantitative fundus autofluorescence as a secondary outcome of a randomized clinical trial. Sci Rep. 2021;11(1):12685.doi: 10.1038/s41598-021-92309-6. PubMed PMID: 34135449. PubMed Central PMCID: PMC8209039.

8. Goh JHL, Lim ZW, Fang X, Anees A, Nusinovici S, Rim TH, et al. Artificial Intelligence for Cataract Detection and Management. Asia Pac J Ophthalmol (Phila). 2020;9(2):88–95.doi: 10.1097/01.Apo.0000656988.16221.04. PubMed PMID: 32349116.

9. Spooner K, Phan L, Cozzi M, Hong T, Staurenghi G, Chu E, et al. Comparison between two multimodal imaging platforms: Nidek Mirante and Heidelberg Spectralis. Graefes Arch Clin Exp Ophthalmol. 2021;259(7):1791–802.doi: 10.1007/s00417-020-05050-7. PubMed PMID: 33409677.

10. Arrigo A, Teussink M, Aragona E, Bandello F, Battaglia Parodi M. MultiColor imaging to detect different subtypes of retinal microaneurysms in diabetic retinopathy. Eye (Lond). 2021;35(1):277–81.doi: 10.1038/s41433-020-0811-6. PubMed PMID: 32066896. PubMed Central PMCID: PMC7852576.

11. Kim YH, Ahn J, Kim KE. Multicolor Imaging for Detection of Retinal Nerve Fiber Layer Defect in Myopic Eyes With Glaucoma. Am J Ophthalmol. 2022;234:147–55.doi: 10.1016/j.ajo.2021.07.022. PubMed PMID: 34314686.

12. Khan Z, Iqbal Z, Iqbal S. Evaluation of the Clinical Application of Multi-Color Optical Coherence Tomography as a Diagnostic Tool for Different Retinal Pathologies. Pakistan Journal of Medical and Health Sciences. 2022;16:1373–5.doi: 10.53350/pjmhs221611373.

13. Yang Y, Jiang Z, Yang C, Xia Z, Liu F. Improved retinex image enhancement algorithm based on bilateral filtering. 4th International Conference on Mechatronics, Materials, Chemistry and Computer Engineering (ICMMCCE 2015); 2015.

14. Land EH. The retinex theory of color vision. Proceedings of the Royal Institution of Great Britain; 1974.

15. Reza AM. Realization of the Contrast Limited Adaptive Histogram Equalization (CLAHE) for Real-Time Image Enhancement. Journal of VLSI signal processing systems for signal, image and video technology. 2004;38(1):35–44.doi: 10.1023/B:VLSI.0000028532.53893.82.

16. Patro S, Sahu KK. Normalization: A preprocessing stage. arXiv preprint 1503.06462. 2015.

17. Simonyan K, Zisserman A. Very deep convolutional networks for large-scale image recognition. arXiv preprint 1409.1556. 2014.

18. Yan Y, Kawahara J, Hamarneh G. Melanoma Recognition via Visual Attention. 2019; Cham.

19. Ilse M, Tomczak J, Welling M. Attention-based deep multiple instance learning. Paper presented at: International conference on machine learning 2018.

20. Woo S, Park J, Lee J-Y, Kweon IS. Cbam: Convolutional block attention module. Paper presented at: Proceedings of the European conference on computer vision (ECCV) 2018.

21. Ekweremadu E, Okoloagu N, Balantine E. The reliability of light projection test in preoperative assessment of cataract patients. Journal of Biomedical Sciences In AFRICA. 2019;16(1&2):20–7.

22. Reid O, Maberley DA, Hollands H. Comparison of the potential acuity meter and the visometer in cataract patients. Eye (Lond). 2007;21(2):195–9.doi: 10.1038/sj.eye.6702165. PubMed PMID: 16273082.

23. Mackenbrock LHB, Łabuz G, Yildirim TM, Auffarth GU, Khoramnia R. Automatic Quantitative Assessment of Lens Opacities Using Two Anterior Segment Imaging Techniques: Correlation with Functional and Surgical Metrics. Diagnostics (Basel). 2022;12(10).doi: 10.3390/diagnostics12102406. PubMed PMID: 36292095. PubMed Central PMCID: PMC9600551.

24. Faria-Correia F, Lopes B, Monteiro T, Franqueira N, Ambrósio R, Jr. Scheimpflug lens densitometry and ocular wavefront aberrations in patients with mild nuclear cataract. J Cataract Refract Surg. 2016;42(3):405–11.doi: 10.1016/j.jcrs.2015.10.069. PubMed PMID: 27063521.

25. Faria-Correia F, Ramos I, Lopes B, Monteiro T, Franqueira N, Ambrósio R, Jr. Comparison of Dysfunctional Lens Index and Scheimpflug Lens Densitometry in the Evaluation of Age-Related Nuclear Cataracts. J Refract Surg. 2016;32(4):244–8.doi: 10.3928/1081597x-20160209-01. PubMed PMID: 27070231.

26. Shih KC, Hung KW, Lau KP, Yip WM, Lee A, Fong A, et al. Deep Learning Automated Diagnosis and Grading of Cataracts using Colour Fundus Images: The Fundus Cataract-AI Project. Investigative Ophthalmology & Visual Science. 2024;65(7):5950-.

27. Okuno T, Kojima M, Yamaguchi-Sekino S, Ishiba Y, Suzuki Y, Sliney DH. Cataract Formation by Near-infrared Radiation in Rabbits. Photochemistry and Photobiology. 2021;97(2):372–6.doi: 10.1111/php.13342.

28. Gorgels TG, van Norren D. Ultraviolet and green light cause different types of damage in rat retina. Invest Ophthalmol Vis Sci. 1995;36(5):851-63. PubMed PMID: 7706033.

